# 216 GHz Electron Paramagnetic Resonance of Mycobacterium Tuberculosis Catalase-Peroxidase: The Role of the Arg418 Residue

**DOI:** 10.1101/246249

**Authors:** Matt Bawn, Jurek Krzystek, Richard Magliozzo

**Author notes:** To whom correspondence should be addressed: Richard Magliozzo, ^1^Department of Chemistry, Brooklyn College of the City University of New York, Brooklyn, New York 11210.

## Abstract

The catalase-peroxidase protein from *Mycobacterium tuberculosis* contains a variety of unique structural features including a covalently-linked three amino acid adduct capable of hosting a tyrosine-based radical. Previous work has demonstrated that the Arg418 residue is essential for the catalse but not the peroxidase activity of the protein and crystallography has indicated the residue to be capable of adopting two conformations relative to the adduct-radical. In the present work the WT and Arg418Leu mutant proteins were investigated using high-field electron magnetic resonance spectroscopy. Different sets of g-values were found for each protein indicating different paramagnetic environments. Quantum chemical calculations of model structures were undertaken to elucidate the geometrical environment of the radical. It is proposed that the two sets of g-values correspond to the two conformations of the Arg418 residue. The implications for the catalytic mechanism are discussed.

## INTRODUCTION

*Mycobacterium tuberculosis* catalase-peroxidase (Mtb KatG) is a dual-function heme containing enzyme that also contains a unique covalently-linked three amino acid adduct consisting of residues Trp107, Tyr229 and Met255 (MYW-adduct). The adduct has previously been shown to be necessary for the catalase but not the peroxidase activity of the protein. The adduct is capable of hosting a tyrosine-based radical as part of the catalase reaction, other work□^1^□ has revealed a critical role for the Arg418 residue in this reaction. The residue has been seen to be able to adopt two conformations relative to the MYW-adduct (vicinal and distal) as revealed by x-ray crystallography^2□^ and shown in **Figure 1**. Electron Paramagnetic Resonance (EPR) spectroscopy is a technique capable of probing the electronic environment of amino acid radicals and has been used to characterize Tyr radicals in a variety of enzymes. Previous high-field EPR spectroscopy undertaken by the group at D-band (130 GHz) of wild-type (WT) Mtb KatG suggested the presence of two MYW-adduct radical environments characterized by two distinct g_x_ values^3□^. To determine whether these two environments correspond to the two conformations of the Arg418 residue EPR has again been applied but this time to the Arg418Leu mutant and WT Mtb KatG proteins to produce the distal and vicinal conformations respectively.

**Figure 1.**
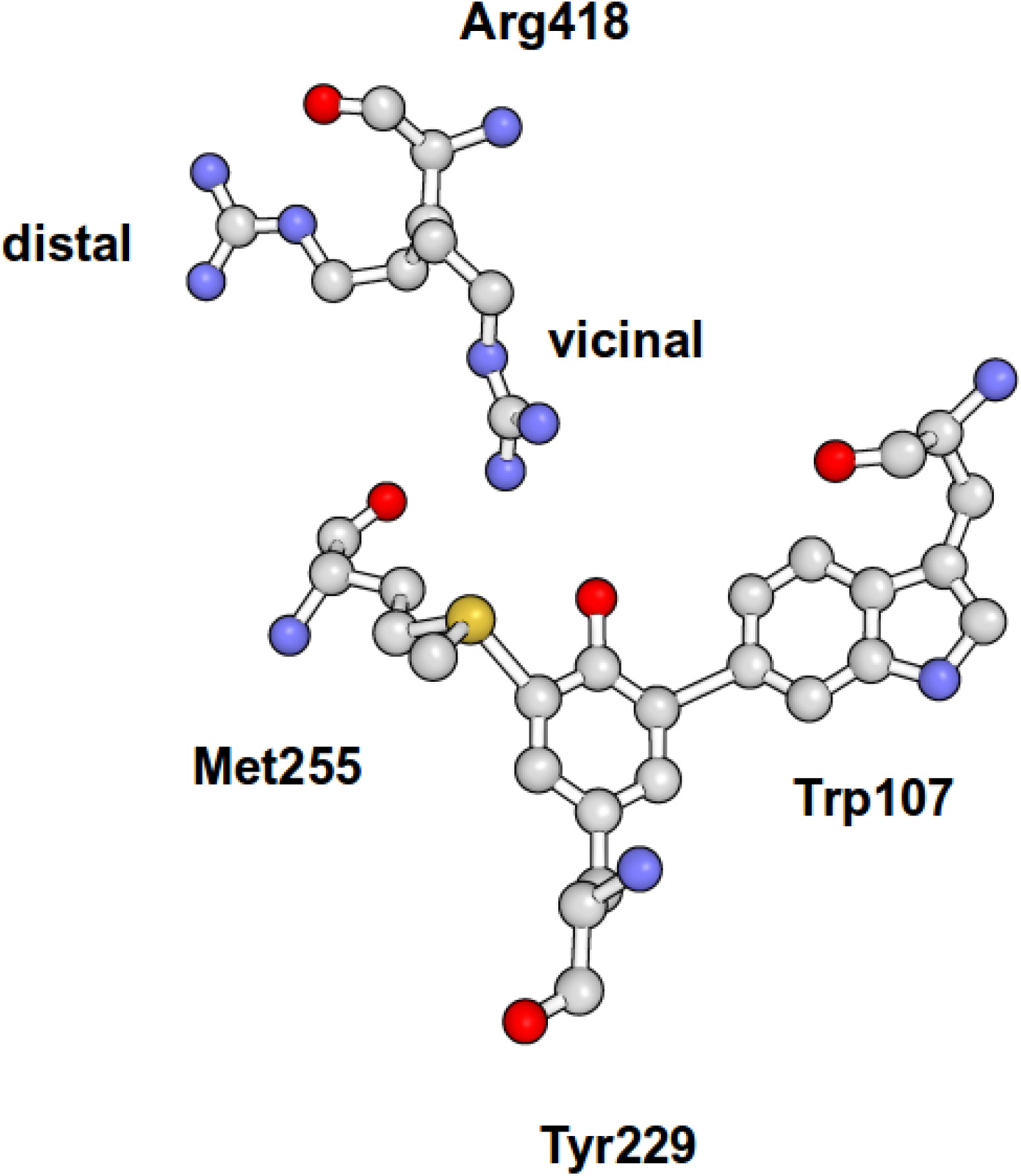
MYW-adduct and Arg418 resides from Mtb KatG [PDB:2CCA]. Arg418 is shown in vicinal and distal conformations.

## EXPERIMENTAL

*Sample Preparation:* WT and Arg418Leu KatG were grown and purified as described previously^1□^. Protein was reacted against 8000-fold excess hydrogen peroxide at pH 7.5 and manually freeze-quenched (freezing time ~ 2 s) in liquid ethane. Excess liquid ethane was removed and the samples transferred at low-temperature to Teflon EPR cups. Samples were stored in liquid nitrogen at 77K when not in use.

*EPR:* Field-modulated continuous-wave EPR spectra were measured at 216 GHz. An atomic hydrogen standard was included during measurement for accurate field determination^4□^.

## RESULTS

*HF-EPR:* The 216 GHz EPR and simulated spectra measured at 15 and 60 K of WT and Arg418Leu KatG respectively are shown in **Figure 2**. In the case of the WT sample there was a large source of Mn(II) contamination that is commonly detected by HF EPR and becomes more apparent at higher temperatures. Temperatures were chosen in order to enhance the derivative nature of the spectra and facilitate simulation. The simulations clearly show g-values for each protein of g_x_ = 2.0064 g_y_ = 2.0038 g_z_ = 2.0024 for the WT and g_x_ = 2.0054 g_y_ = 2.0033 g_z_ = 2.0019 Arg418Leu. The contributions to both spectra from Mn(II) are also shown in the figure.

**Figure 2.**
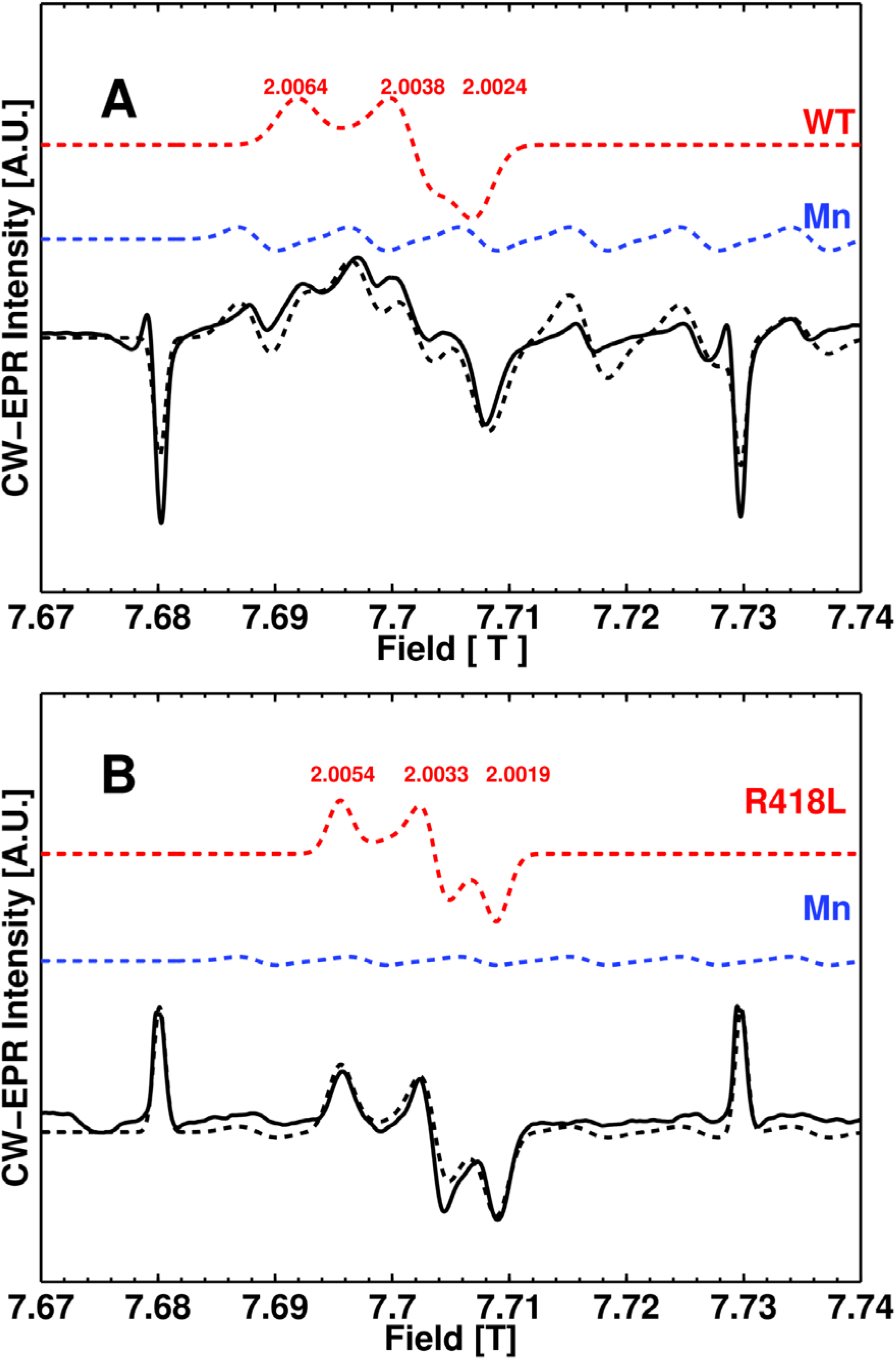
216 GHz continuous-wave EPR spectra (solid) and simulations (dashed) measured at 15 Kand 60 K respectively of WT (A) and Arg418Leu mutant Mtb KatG (B). Simulation parameters for WT were: g_x_ = 2.0064 g_y_ = 2.0038 g_z_ = 2.0024 whilst those for Arg418Leu were: g_x_ = 2.0054 g_y_ = 2.0033 g_z_ = 2.0019. The contributions to the simulation from Mn(II) and KatG species are indicated in blue and red respectively.

**Figure 3.**
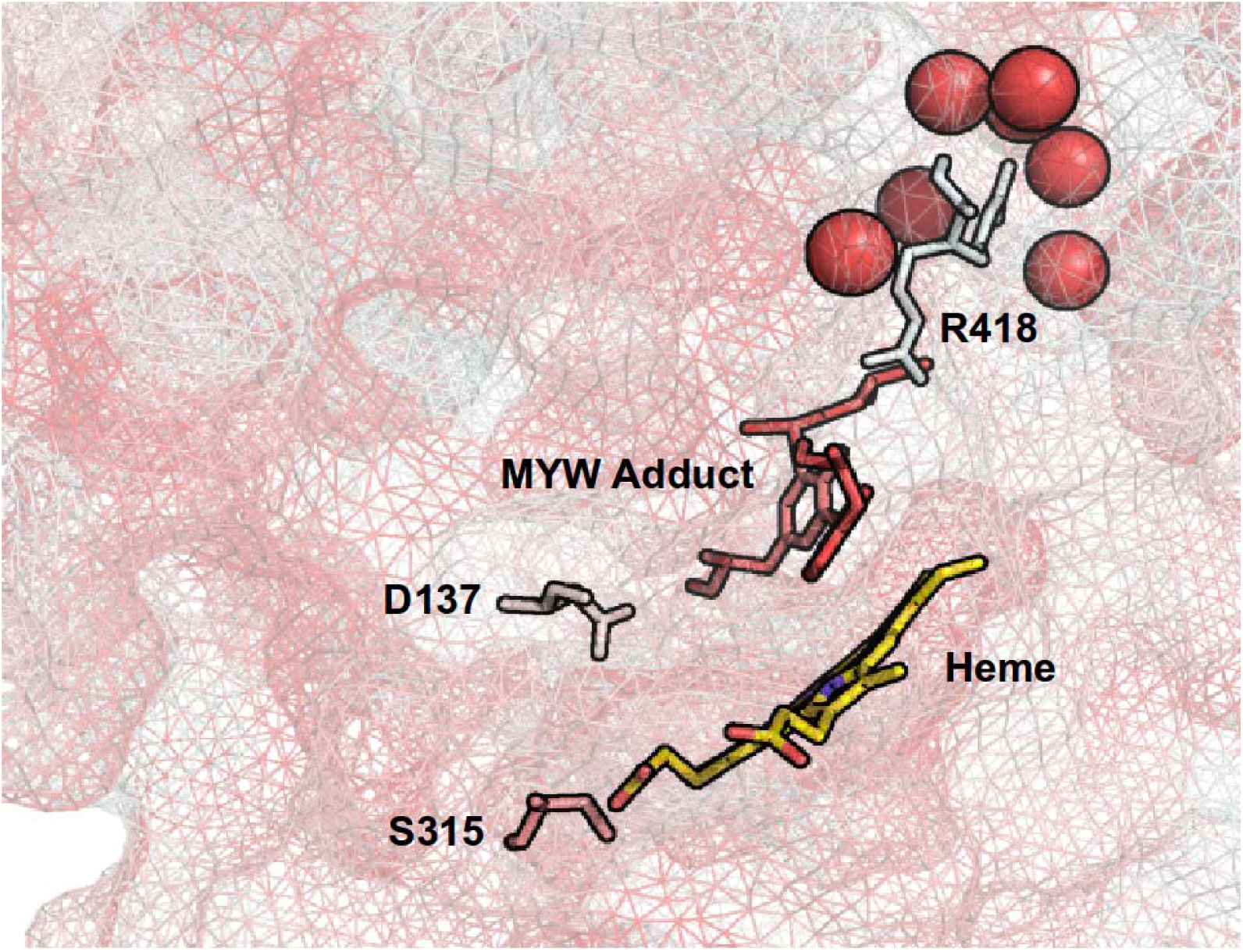
Hydrophobicity of heme binding pocket environment of KatG. [PDB:2CCA] The strength of hydrophobicity is indicated by the degree of redness of the wire-frame representation. Crystollagraphic water oxygen atoms within 5 nm of the R418 residue are shown as red spheres.

## DISSCUSION

Previous spectra measured at 130 GHz and 7K of the WT protein at pH 8.5 also revealed a rhombic gtensor with dual g_x_ values (2.00550 and 2.00606), g_y_ = 2.00344 and g_z_ = 2.00186. Theses are in reasonable agreement with the g-values obtained in this study for the WT and Arg418Leu mutant protein. It must also be remembered however, that these previous results were performed on slightly different samples. The main differences to be taken into consideration should be the freezing time (35 ms RFQ for the D-band vs 2 s MFQ in the present study) and the pH (pH 8.5 for D-band and pH 7.5 inthe present work). Indeed in another heme protein in which an Arg residue has been shown to be able to “swing” into a vicinal and distal position, pH has been indicated to be a factor in the conformation.

The results of the HF-EPR clearly show the presence of two differing conformations of the paramagnetic environment in the WT and ARG418Leu proteins that may be explained by the ARG418 residue being in the vicinal and distal conformations. The presence of a bi-conformational ARG residue has also been seen in the crystal structure of versatile peroxidase^11□^. The Trp164Tyr mutant of the protein however, seemed to stabilize only one of the ARG conformations.

DFT calculations of the g-tensor for the MYW radical in katG with the R418 residue in both conformations indicate that the g_x_-value for the vicinal ARG should be lower than for the distal position. This is opposite to what is observed experimentally in the case of the WT and R418L HF-EPR results described in this work.

## REFERENCES

Zhao, X. et al. Specific function of the Met-Tyr-Trp adduct radical and residues Arg-418 and Asp-137 in the atypical catalase reaction of catalase-peroxidase KatG. J. Biol. Chem. 287, 37057–65 (2012).

Zhao, X. et al. Hydrogen peroxide-mediated isoniazid activation catalyzed by Mycobacterium tuberculosis catalase-peroxidase (KatG) and its S315T mutant. Biochemistry45, 4131–4140 (2006).

Suarez, J. et al. An Oxyferrous Heme / Protein-based Radical Intermediate Is Catalytically Competent in the Catalase Reaction of Mycobacterium tuberculosis Catalase-Peroxidase (KatG) ^*^ □. J. Biol. Chem. 284, 7017–7029 (2009).

Stoll, S., Ozarowski, A., Britt, R. D. & Angerhofer, A. Atomic hydrogen as high-precision field standard for high-field EPR. J. Magn. Reson. 207, 158–63 (2010).

Becke, A. D. Density-Functional Thermochemistry. 3. The Role of Exact Exchange. J. Chem. Phys. 98, 5648–5652 (1993).

Lee, C. T., Yang, W. T. & Parr, R. G. Development of the Colle-Salvetti Correlation-Energy Formula into a Functional of the Electron-Density. Phys. Rev. B37, 785–789 (1988).

Stephens, P. J., Devlin, F. J., Chabalowski, C. F. & Frisch, M. J. Ab Initio Calculation of Vibrational Absorption and Circular Dichroism Spectra Using Density Functional Force Fields. J. Phys. Chem. 98, 11623–11627 (1994).

Hehre, W. J. Self—Consistent Molecular Orbital Methods. XII. Further Extensions of Gaussian— Type Basis Sets for Use in Molecular Orbital Studies of Organic Molecules. J. Chem. Phys. 56, 2257 (1972).

Francl, M. M. Self-consistent molecular orbital methods. XXIII. A polarization-type basis set for second-row elements. J. Chem. Phys. 77, 3654 (1982).

Schafer, A., Huber, C. & Ahlrichs, R. Fully Optimized Contracted Gaussian-Basis Sets of Triple Zeta Valence Quality for Atoms Li to Kr. J. Chem. Phys. 100, 5829–5835 (1994).

Bernini, C. et al. EPR parameters of amino acid radicals in P. eryngii versatile peroxidase and its W164Y variant computed at the QM/MM level. Phys. Chem. Chem. Phys. 13, 5078–98 (2011).

